# Temperature-responsive structural reversibility of FGF21 and structure-based design of its variant with enhanced potency

**DOI:** 10.1101/2021.11.16.468794

**Authors:** Ye-Eun Jung, Kyeong Won Lee, Jae Hyun Cho, Da-Woon Bae, Bo-Gyeong Jeong, Yeon-Ju Jung, Young Jun An, Kyungchan Kim, Ga Seul Lee, Lin-Woo Kang, Jeong Hee Moon, Jung-Hyun Lee, Eun-Kyoung Kim, Hyung-Soon Yim, Sun-Shin Cha

## Abstract

Fibroblast growth factor 21 (FGF21) has pharmaceutical potential against obesity-related metabolic disorders, including non-alcoholic fatty liver disease. Since thermal stability is a desirable factor for therapeutic proteins, we investigated the thermal behavior of human FGF21. FGF21 remained soluble after heating; thus, we examined its temperature-induced structural changes using circular dichroism (CD). FGF21 showed inter-convertible temperature-specific CD spectra. The CD spectrum at 100 °C returned to that at 20 °C when the heated FGF21 solution was cooled. Through loop swapping, the connecting loop between β10 and β12 in FGF21 was revealed to be associated with the unique thermal behavior of FGF21. According to *in vitro* cell-based assays and model high-fat diet (HFD)-induced obesity studies, heated FGF21 maintained biological activities that were comparable to those of non-heated and commercial FGF21s. Based on sequence comparison and structural analysis, five point-mutations were introduced into FGF21. Compared with the wild type, the heated FGF21 variant displayed improved therapeutic potential in terms of body weight loss, the levels of hepatic triglycerides and lipids, and the degree of vacuolization of liver in HFD-fed mice.

## Introduction

Fibroblast growth factors (FGFs) regulate various developmental and metabolic processes, including cell proliferation, differentiation, angiogenesis, wound healing, nerve regeneration, chronic inflammation, and cancer growth (1, 2). The FGF family consists of 22 members that share the β-trefoil fold despite their relatively low sequence identities (13–71%) (3). The β-trefoil fold contains 12 β-strands that form 6 two-stranded β-hairpins. The first β-12 strands, while the others comprise two successive β-strand pairs (i.e. β2-β3-β4-β5-β6-β7-β-8-β9, and β10 -β11). Hairpins are arranged in a pseudo three-fold symmetry: the first, third, and fifth β-hairpins form a barrel that is covered by a triangular cap consisting of the second, fourth, and sixth β-hairpins (3, 4).

FGFs are divided into FGF1, FGF4, FGF7, FGF8, FGF9, FGF11, and FGF19 subfamilies depending on their sequence similarities: subfamilies. Except for the intracellular FGF11 subfamily, FGFs form complexes with FGF receptors (FGFRs) on the cell membrane, which are generally composed of three immunoglobulin (Ig)-like extracellular domains, a transmembrane domain, and a cytoplasmic tyrosine kinase domain (2,3,5). FGFs bind to the interface between the second and third Ig-like ectodomains to induce receptor dimerization, leading to the phosphorylation of tyrosine residues in the cytoplasmic domain to stimulate signaling pathways (1,2,6).

The canonical FGFs (FGF1, FGF4, FGF7, FGF8, and FGF9 subfamilies) have a positively charged sector clustered by lysine and arginine residues and thus display avidity for negatively charged heparin/heparan sulfate proteoglycans (HSPGs) present on the cell surface or in the extracellular matrix (7). Therefore, canonical FGFs form ternary complexes with HSPG and FGFR, and act in the vicinity of cells as paracrine and/or autocrine factors. Conversely, FGF19 subfamily members (FGF19, FGF21, and FGF23) have no apparent HSPG-binding site on their surfaces and thus perform their physiological roles in an endocrine manner (8). They are released from the extracellular matrix and reach remote target organs through the bloodstream (9). A remarkable structural feature of FGF19 subfamily members is the presence of an additional C-terminal loop protruding from the β-trefoil core structure. The long C-terminal loop comprising ∼30 residues tightly binds to α-Klotho (10) or β-Klotho (11, 12), known as the co-receptors of FGF23 or FGF19 and FGF21, respectively. The truncation of the C-terminal tail or the absence of klotho causes severe defects in FGF signaling, demonstrating the critical role of the interaction between the C-terminal loop and klotho (10,13,14).

Since klothos are membrane-bound proteins expressed in certain organs, their presence determines the tissue selectivity of FGF19 subfamily members that form the ternary FGF-FGFR-klotho complex (15, 16). β-Klotho is mainly distributed in adipose tissues, the liver, and the pancreas, indicating that those are the target sites of FGF21; in fact, FGF21 has been implicated in glucose and lipid metabolism (16, 17). Administration of FGF21 to rodents or non-human primates causes a reduction in fat mass and body weight, as well as levels of circulating glucose and triglycerides (TG), and improves insulin sensitivity and energy metabolism (18–20). Therefore, FGF21 shows therapeutic potential to treat obesity-related metabolic complications, including hyperglycemia, insulin resistance, and non-alcoholic fatty liver disease (NAFLD) (21–23).

There are several hurdles that should be tackled to develop FGF21 for clinical application. For example, the C-terminal loop of FGF21 is essential for its activity due to strong interactions with β-Klotho, but it is highly susceptible to proteolytic attack (24). Proteases in expression hosts can inactivate FGF21 by breaking this loop during purification processes. Heating that precipitates heat-labile host proteases can minimize the proteolytic degradation of the C-terminal loop, if FGF21 endures heat treatment. In this study, we discovered the unreported thermal behavior of human FGF21 and employed this property for efficient purification of active human FGF21 (hereafter, FGF21). Additionally, structure-based designing was performed to develop FGF21 variants with point mutations in the C-terminal region interacting with β-Klotho, which displayed enhanced potency for *in vitro* and *in vivo* studies compared with the wild type.

## Results and discussion

### Temperature-responsive structural reversibility of FGF21

To investigate how susceptible FGF21 is to thermal denaturation, FGF21-containing solutions were incubated at high temperatures for 10 min. When proteins are denatured, they form coarse aggregate particles that scatter visible light to make solutions turbid. There was no change in turbidity of FGF21 solutions stored at 20 °C after incubation at 40, 60, 80, and 100 °C, when observed by naked eyes, indicating the absence of large protein aggregates (Fig. S1). In contrast, FGF2 solutions irreversibly became turbid after heating; turbidity was maintained regardless of storing temperatures (Fig. S1). To confirm that FGF21 is resistant to thermal denaturation, the heat-treated FGF21 solutions were centrifuged to remove protein aggregates, and then supernatants were analyzed using SDS-PAGE. As shown in Figure 1*A*, the amount of FGF21 in the solutions was nearly identical before and after heat treatment, indicating that no protein aggregates were formed due to thermal denaturation. However, FGF2 was dramatically reduced in supernatants after incubation at high temperatures (Fig. 1*A*).

**Figure 1.**
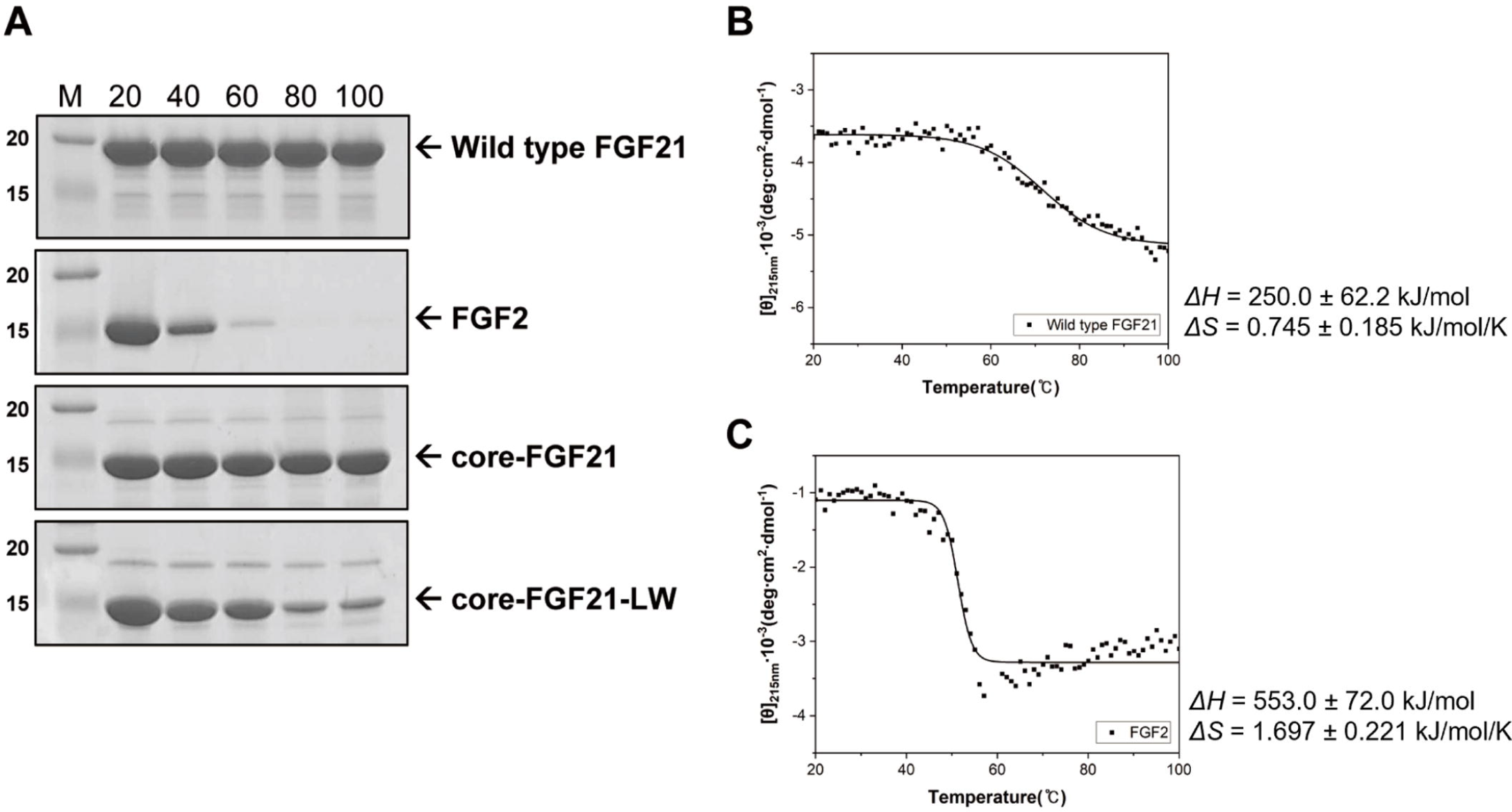
Monitoring the thermal denaturation of FGF21. *(A)* SDS-PAGE. M represents protein markers whose sizes are given in kDa on the left. Numbers in the top are incubation temperatures (°C). Arrows point out the position of target proteins. Thermal denaturation curves of FGF21 *(B)* and FGF2 *(C)*. Spectral curves were fitted using the molar ellipticity [θ] values shown as dots.

Thermal denaturation curve is a good indicator for assessing the thermal stability of proteins. To assess the thermal denaturation of FGF21 using CD spectroscopy, spectral changes were monitored at 215 nm at a temperature range of 20–100 °C (25, 26). The FGF21 spectra displayed gradual changes as a function of temperature, which contrasted with the steep transition slope of the FGF2 spectra (Figs. 1, *B* and *C*). The magnitude of enthalpy and entropy changes was smaller in FGF21 than FGF2 (Figs. 1, *B* and *C*), and the *T*_m_ values were estimated to be 62.26 ± 0.6306 °C and 52.69 ± 0.1991 °C for FGF21 and FGF2, respectively.

For further characterization of the temperature responsiveness of FGF21, we examined the secondary and tertiary structural states of FGF21 as the temperature increased from 20 °C to 100 °C by measuring far-UV (190–260 nm) and near-UV (240–340 nm) CD spectra, respectively (27). In both CD spectra, temperature-induced spectral changes are sequential; for example, the spectrum at 60 °C is located between those at 40 °C and 80 °C (Figs. 2, *A* and *B*). The spectral feature of FGF21 accompanied by temperature change encouraged us to measure the CD spectra while cooling the heated FGF21 solutions. The CD spectra of FGF21 at 20 °C were virtually identical to that of FGF21 that was cooled to 20 °C after heating at 100 °C (Figs. 2, *C* and *D*), indicating that structural changes of FGF21 induced by heating can be restored by cooling (Fig. 2*E*). Consequently, FGF21 is highly likely to have temperature-responsive structural reversibility.

**Figure 2.**
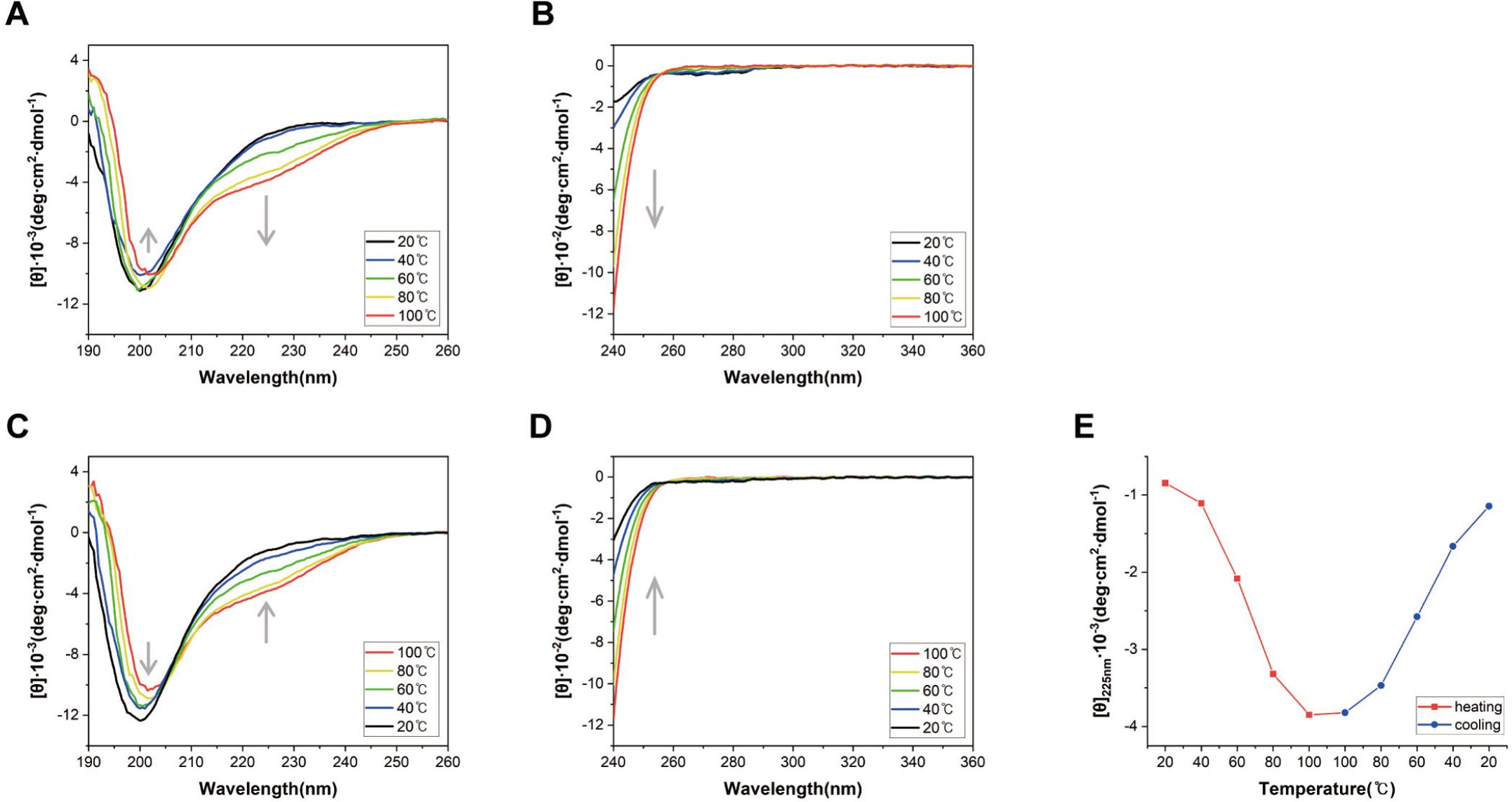
Temperature-dependent CD spectra of the wild type FGF21. Far-UV CD spectra upon heating *(A)* and cooling *(C)*, and near-UV CD spectra upon heating *(B)* and cooling *(D)*. Profiles at different temperatures are represented using different colored lines as shown in the right box of the Figure. Gray arrows indicate the directions of spectral changes during heating and cooling. *(E)* The molar ellipticity [θ] values as temperature increases and decreases are colored in red and blue, respectively. For this Figure, [θ] of FGF21 at 225 nm was used since far-UV CD spectral changes were dramatic at this wavelength.

### The unique loop region associated with the heat response of FGF21

The N- and C-terminal loops extending out from the β-trefoil core structure of FGF21 (core-FGF21) show large sequential disparities when compared with the corresponding regions of other FGFs (Fig. S2). Therefore, we constructed a deletion mutant without the N-terminal loop (Δ174–209) to investigate whether both distinct terminal loops are related to the remarkable response of FGF21 to heating. However, similar to the wild type FGF21, the deletion mutant remained soluble after incubation at high temperatures (Fig. 1*A*), indicating that core-FGF21 is responsible for the thermal behavior of FGF21.

-trefoil core structure with disparities in loop regions connecting β- strands (3, 4). Among connecting loops, the 10-12 loop (residues His145-Pro161) of FGF21 is outstanding in that it has a unique amino acid composition. First, the sequence identities of the 10- 12 loop in FGF21 to corresponding loops in other FGFs lie in the range of 5–25%, whereas the entire FGF21 shows 14–36% sequence identities to other FGFs. Second, the β10- β12 loop of FGF21 contains 5 proline residues, constituting 29.4% of the loop, which contrasts with the fact that most FGFs have 0 or 1 proline residue in the corresponding loop (Fig. S2). Proline residues in loop regions have been reported to contribute to the thermal stability of proteins (28, 29).

Based on these analyses, we tested the contribution of the unique β10-β12 loop to the thermal response of FGF21 through loop swapping. In terms of sequence identity, the β10- β12 loop of FGF23 is most similar to that of FGF21 (Fig. S2), implying that loop swapping with the loop of FGF23 would have minimal structural effect (Fig. 3). Therefore, the β10-β12 loop of core-FGF21 was replaced by the corresponding loop (residues 137–154) of FGF23 to make a loop-swapped mutant (core-FGF21-LW). As shown in Figure 1*A*, the amount of core-FGF21 in solutions was nearly identical, regardless of heat treatment. However, core-FGF21-LW in supernatants was reduced by more than 70% after 10 min incubations at 80 °C and 100 °C (Figs. 1*A* and S1). The formation of protein aggregates was monitored by measuring optical density since aggregates scatter visible light strongly due to their large sizes (30). A solution containing core-FGF21-LW displayed the initial optical density at 600 nm (OD600) of 0.005 when incubated at 75 °C. However, over the duration of the treatment, the OD600 value of the solution reached 1.086, indicating that core-FGF21-LW formed aggregates. In contrast, the OD600 value of a solution containing core-FGF21 remained constant, which led us to assume that the β10-β12 loop is related to the heat response of FGF21 (Fig. S3). Similar to the wild type, the loop-swapped FGF21 mutant with the N- and C-terminal loops (FGF21-LW) activated FGF receptor 1c (FGFR1c) (Fig. 3*A*), and its CD spectra in the far-UV region (190–260 nm) was similar to that of the wild type (Fig. 3*B*). Taken together, the loop replacement disturbed only the thermal behavior without affecting the structural and functional properties of FGF21.

**Figure 3.**
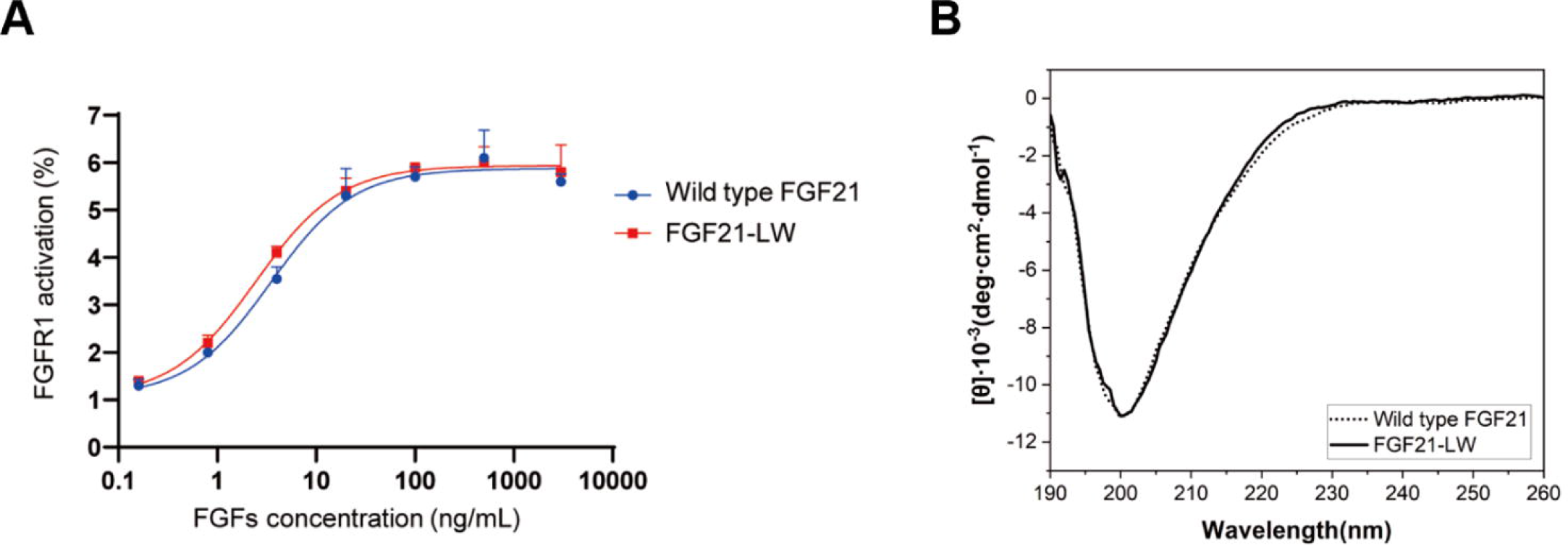
The maintenance of structural and functional properties of the loop-swapped FGF mutant. *(A)* Cell-based activity tests of FGF21-LW. Modified HEK293 cells for reporting FGFR1c activity were treated with the wild type FGF21 or FGF21-LW at the indicated concentration for 6 h. Cell lysates were prepared for detection of luciferase activity. The levels of FGFR1c activity in non-treated cells were set to 1, and the other values were calculated relative to this value (n = 2, average ± SD). *(B)* Comparison of Far-UV CD spectra between the wild type FGF21 and FGF21-LW.

### Heat treatment purification of active human FGF21

*E. coli* is a mesophilic organism frequently used to produce recombinant proteins. Since most *E. coli* proteins are heat-labile, incubation at high temperatures is an efficient strategy for the purification of thermostable proteins produced in *E. coli*. As described in the “Experimental procedures” section, heat treatment at 100 °C was implemented to purify FGF21. In our construct design, FGF21 was expressed with the N-terminal thioredoxin (Trx) tag. Since Trx is heat stable, the Trx-FGF21 fusion protein was incubated at 100 °C for 10 min (31), which separated the Trx-FGF21 protein from heat-labile *E. coli* proteins that precipitated at high temperatures (Fig. S4) (32). Subsequently, the Trx tag was removed by the cleavage of the linker between Trx and FGF21 to secure only FGF21 (33). According to dynamic light scattering (DLS) analysis, the size of FGF21 purified through heating treatment (htFGF21) was highly similar to that of FGF21 purified without heating (nFGF21) (Fig. S5). This strongly suggests that heat treatment has little influence on the quality of FGF21 (34).

Our CD and DLS analyses revealed that heat treatment had little effect on the structure of FGF21 (Figs. 2 and S5). Therefore, we investigated whether heat treatment influences the activity of FGF21 by comparing FGFR1c activation levels of nFGF21, htFGF21, and commercial FGF21 (cFGF21) using FGF21-responsive reporter cells (modified HEK293 cells) that allow for monitoring FGFR1c activation. Both nFGF21 and htFGF21 displayed little difference in FGFR1c activation levels, which were comparable to the activity of cFGF21 (Fig. 4*A*). Since adipocytes are one of the major targets of FGF21, we also examined the activities of the three FGF21s by detecting the phosphorylation of FGFR substrate 2 (FRS2) and extracellular signal-regulated kinase (ERK) in 3T3L1 adipocytes. The treatment of nFGF21, htFGF21, and cFGF21 led to similar phosphorylation levels of FRS2 and ERK (Figs. 4, *B* and *C*). Furthermore, the levels of glucose uptake by FGF21s were similar in 3T3L1 adipocytes (Fig. 4*D*). These results indicate that FGF21 remains active after heating.

**Figure 4.**
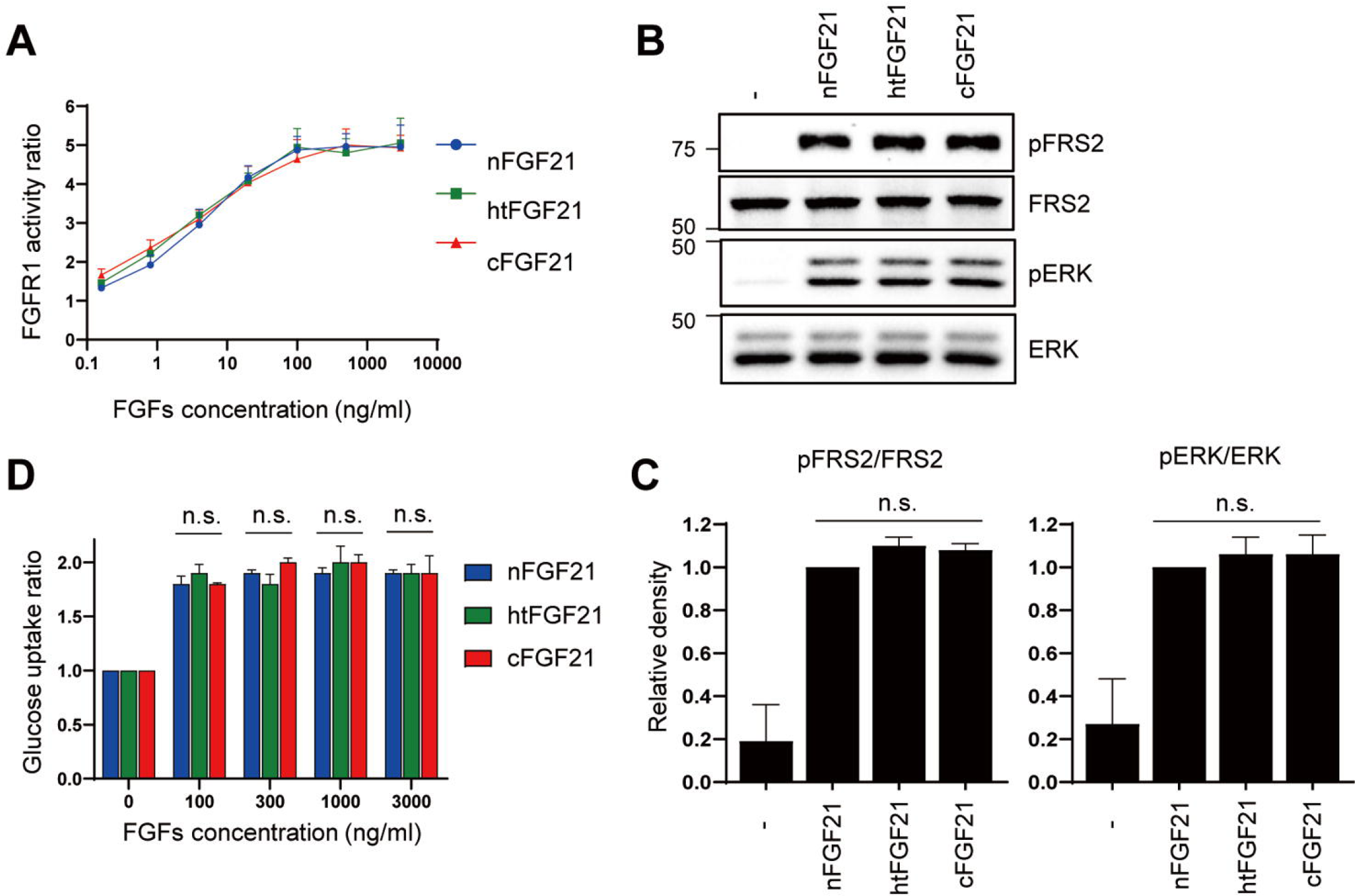
Cell-based activity tests of htFGF21. *(A)* Modified HEK293 cells used for reporting the FGFR1c activity were treated with nFGF21, htFGF21, or cFGF21 at the indicated concentration for 6 h. Cell lysates were prepared for detection of luciferase activity. The levels of FGFR1c activity in non-treated cells were set to 1, and the other values were calculated relative to this value (n = 3, average ± SEM). *(B)* 3T3L1 adipocytes were treated with 100 ng/mL FGF21s for 15 min. Protein samples were prepared for western blotting. *(C)* The band density of nFGF21 treated cells was set to 1, and the other values were calculated relative to this value (n = 3, average ± SEM). *(D)* 3T3L1 adipocytes were treated with FGF21s at the indicated concentration for 6 h, followed by detection of glucose uptake levels. The glucose uptake values of non-treated cells were set to 1, and the other values were calculated relative to this value (n = 3, average ± SEM). The significance was evaluated using the one-way ANOVA, followed by Tukey’s post hoc test. n.s., not significant.

The administration of human or murine FGF21 alleviates the progression of NAFLD in HFD-induced obesity models (35–37). Considering that nFGF21 has traditionally been used in those previous studies, we sought to determine whether htFGF21 was also effective in ameliorating fatty liver. To investigate the effects of htFGF21 on NAFLD, HFD-fed mice were intraperitoneally injected with 1 or 10 mg/kg htFGF21, once daily for 14 days. The injections of either 1 or 10 mg/kg htFGF21 markedly reduced body weight in HFD-fed mice compared with vehicle controls (Figs. 5, *A* and *B*). Significant increases in intrahepatic TG contents were observed in vehicle-treated HFD-fed mice compared with vehicle-treated normal chow diet (NCD)-fed mice; however, the HFD-induced increases in intrahepatic TG dramatically diminished after treatment with 1 or 10 mg/kg htFGF21 (Fig. 5*C*). Histological analyses of liver tissue stained with hematoxylin and eosin (H&E) showed that there was extensive hepatocyte vacuolation in vehicle-treated HFD-fed mice, suggesting intrahepatic fat accumulation (Fig. 5*D*). Conversely, few hepatocellular vacuolations were observed in liver sections of either 1 or 10 mg/kg htFGF21-treated HFD-fed mice, which was comparable with that of vehicle-treated NCD-fed mice (Fig. 5*D*). Moreover, Oil red O staining of liver sections showed that htFGF21 injections dramatically attenuated the formation of intense lipid droplets in HFD-fed mice (Fig. 5*D*). These results suggest that htFGF21 treatment can prevent HFD-induced NAFLD in mice. Collectively, cell-based functional assays and *in vivo* mouse model studies strongly suggest that heating allows for an easy and effective purification of active FGF21.

**Figure 5.**
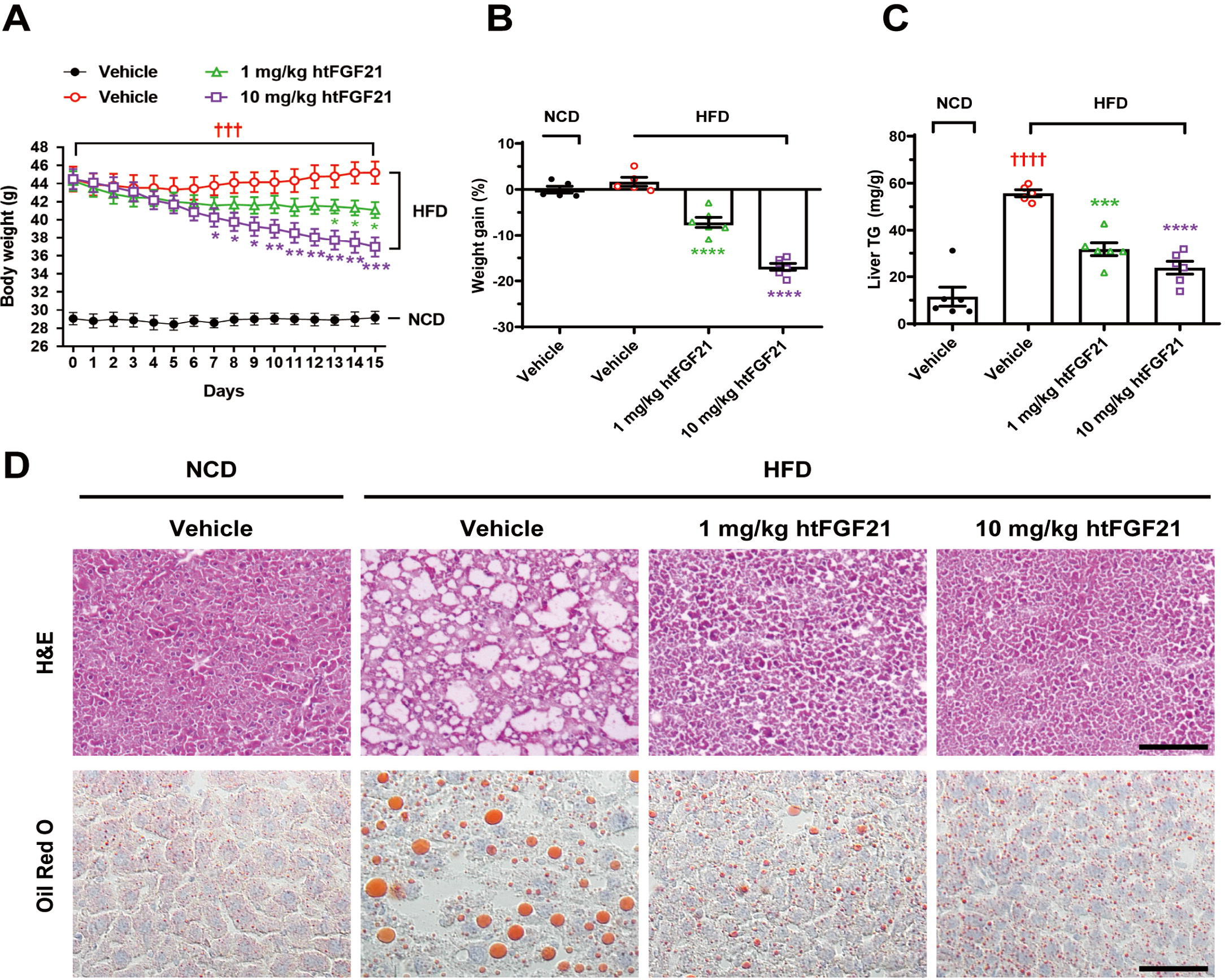
*In vivo* activity tests of htFGF21. HFD-fed mice were injected with htFGF21 intraperitoneally at doses of 0 (vehicle), 1, or 10 mg/kg once daily for 14 days. NCD-fed mice were included as controls and injected intraperitoneally with vehicles. *(A)* Changes in body weight throughout the injection. *(B)* Percent weight gains on the following day after the last injection. *(C)* Effects of the injection on liver TG contents. *(D)* H&E and Oil Red O staining of liver sections showing reversal of vacuolation and lipid accumulation in htFGF21- injected HFD-fed mice. Data are represented as mean ± SEM, calculated using one-way ANOVA followed by Tukey’s multiple comparisons test, * *p* < 0.05, ** *p* < 0.01, *** and ††† *p* < 0.001, **** and †††† *p* < 0.0001. n = 5–6 mice/group. ††† and †††† indicate vehicle- injected NCD-fed mice versus vehicle-injected HFD-fed mice. *, **, ***, and **** indicate vehicle-injected HFD-fed mice versus htFGF21-injected HFD-fed mice. Scale bars: 100 μm (H&E), 50 μm (Oil Red O).

### Structure-based design of a FGF21 mutant with enhanced potency

According to the crystal structure of the C-terminal loop of human FGF21 complexed with β-Klotho (11, 12), residues at positions 201 and 208 are directly involved in interactions with β-Klotho; the side chains of Gln201 and Ala208 fit into a small groove and canyon, respectively, on the surface of β-Klotho (Fig. S6). Sequence comparison shows that Gln201 and Ala208 of human FGF21 are replaced by His201 and Thr208, respectively, in some mammalian FGF21s (Fig. S7). To assess the effects of the two amino acids replacements, we introduced Q201H and A208T mutations into the C-terminal loop of FGF21. The htFGF21 variant with Q201H and A208T mutations (htM2) displayed 2-fold increased potency for FGFR1c activation than the wild type htFGF21 (Fig. 6*A*). In terms of the structural aspect, the C-terminal loop protrudes from the main body of FGF21 and thus is vulnerable to proteolytic degradation. Fibroblast activation protein (FAP) in serum inactivates FGF21 by cleaving the peptide bond between Pro199 and Ser200 in the C-terminal loop, and the P199G mutation is effective to prevent the FAP-mediated cleavage of the loop (24). With a rationale that the enhanced flexibility of the C-terminal loop can have a positive effect on β-Klotho binding (38), we introduced two more glycines to the C-terminal loop (S195G and S200G) in addition to the P199G mutation. In the crystal structure of the C-terminal loop complexed with β-Klotho (11), the side chains of both serine residues are exposed to the solvent without interacting with β-Klotho (Fig. S6). Therefore, the Ser -7 Gly mutations are likely to have no effect on β-Klotho interactions but could increase the loop flexibility. The htFGF21 variant with S195G, P199G, and S200G mutations (htM3), which has three glycine residues in the C-terminal loop, showed 3.65-fold higher potency for FGFR1c activation than the wild type htFGF21 (Fig. 6*A*). Encouraged by the two mutational studies that produced mutant FGF21s with higher FGFR1c-activating functions, we combined the five point-mutations in the C-terminal loop. The htFGF21 variant with S195G, P199G, S200G, Q201H, and A208T mutations (htM5) showed the highest potency for FGFR1c activation; this htM5 mutant showed 5.27-fold higher potency than the wild type htFGF21 (Fig. 6*A*). The activities of htM2, htM3, and htM5 were also confirmed in 3T3L1 adipocytes. Consistent with FGFR1c activation in Figure 6*A*, the three mutants increased the levels of phosphorylated FRS2 and ERK more than the wild type htFGF21 (Figs. 6, *B* and *C*).

**Figure 6.**
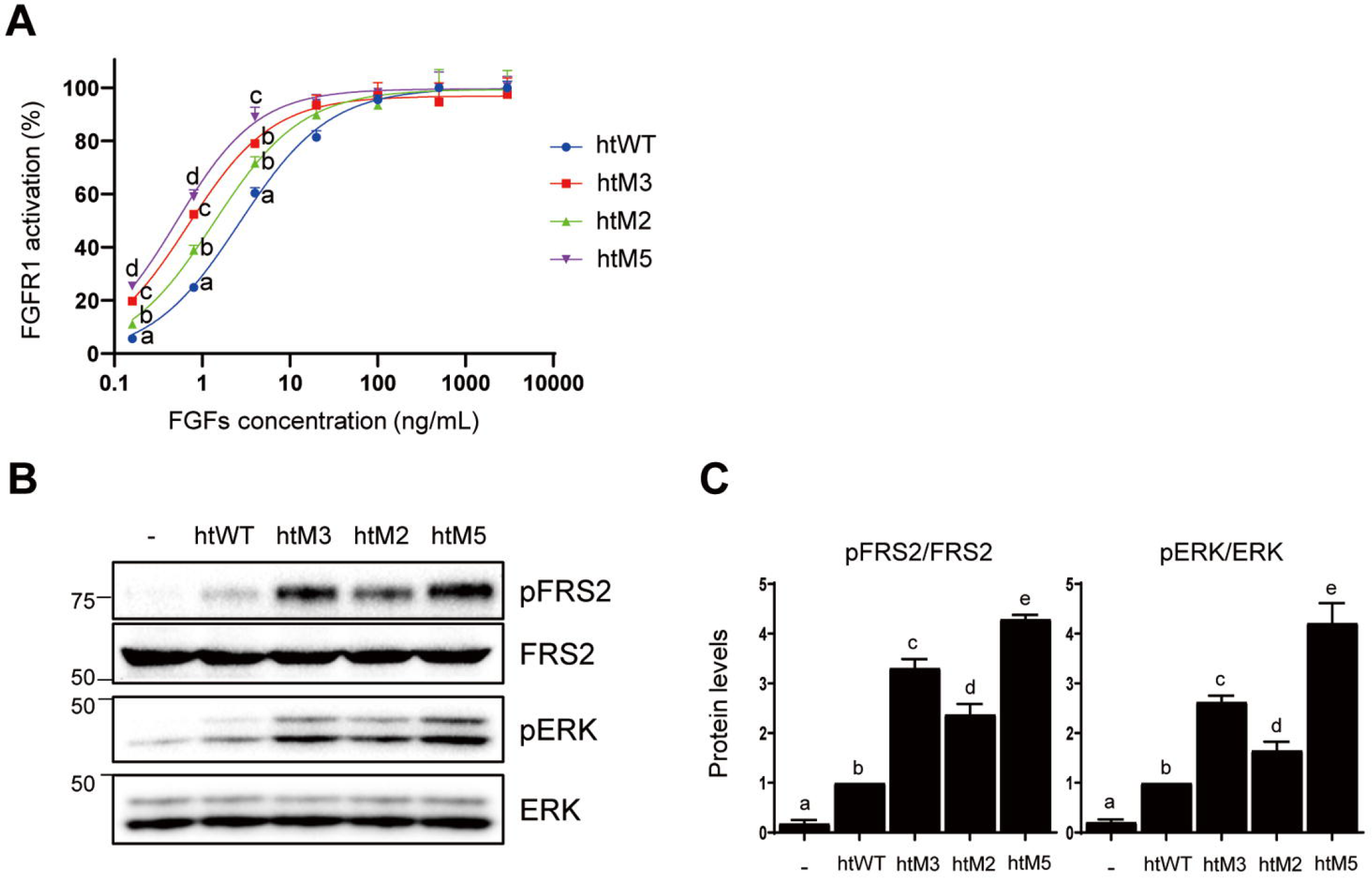
Cell-based activity tests of htM2, htM3, and htM5. *(A)* Modified HEK293 cells used for reporting FGFR1c activity were treated with the wild type htFGF21 (htWT), htM2, htM3, and htM5 at the indicated concentration for 6 h. Cell lysates were prepared for detection of luciferase activity. The levels of FGFR1c activity in non-treated cells were set to 1, and the other values were calculated relative to this value (n = 4, average ± SEM). Non- linear regression was performed using GraphPad Prism 8, and the best-fit non-linear regression curve is depicted. EC_50_s of htWT, htM2, htM3, and htM5 were calculated as 2.74 ± 0.24, 1.37 ± 0.06, 0.75 ± 0.17, and 0.52 ± 0.01 ng/ml (average ± SEM), respectively. The statistical differences at each concentration were evaluated using one-way ANOVA followed by Tukey’s post hoc test and are represented as a, b, c, and d. *p* < 0.05. *(B)* 3T3L1 adipocytes were treated with htWT, htM2, htM3, and htM5 30 ng/mL for 15 min. Protein samples were prepared for western blotting. *(C)* The band density of htWT treated cells was set to 1, and the other values were calculated relative to this value (n = 3, average ± SEM). Statistical significance was evaluated using one-way ANOVA followed by Tukey’s post hoc test and was marked as a, b, c, d, and e. *p* < 0.05.

To compare effects on NAFLD between the wild type and htM5, HFD-fed mice were intraperitoneally injected with 0.1 or 0.5 mg/kg of htFGF21 or htM5, once daily for 14 days. Although the injections of either the 0.1 mg/kg htFGF21 or htM5 decreased body weight compared with vehicle-treated mice, the trend for body weight loss was greater in the htM5 group than in the htFGF21 group (Figs. 7, *A* and *B*). Mice injected with either the 0.5 mg/kg htFGF21 or htM5 treatment showed significant body weight loss compared with vehicle-treated mice, and there was no difference in the extent of weight loss between the htFGF21 and htM5 groups (Figs. 7, *A* and *B*). Either the htFGF21 or htM5 injections substantially reduced hepatic TG content in a dose-dependent manner but htM5 significantly decreased hepatic TG content more than htFGF21 at 0.1 mg/kg (Fig. 7*C*).

**Figure 7.**
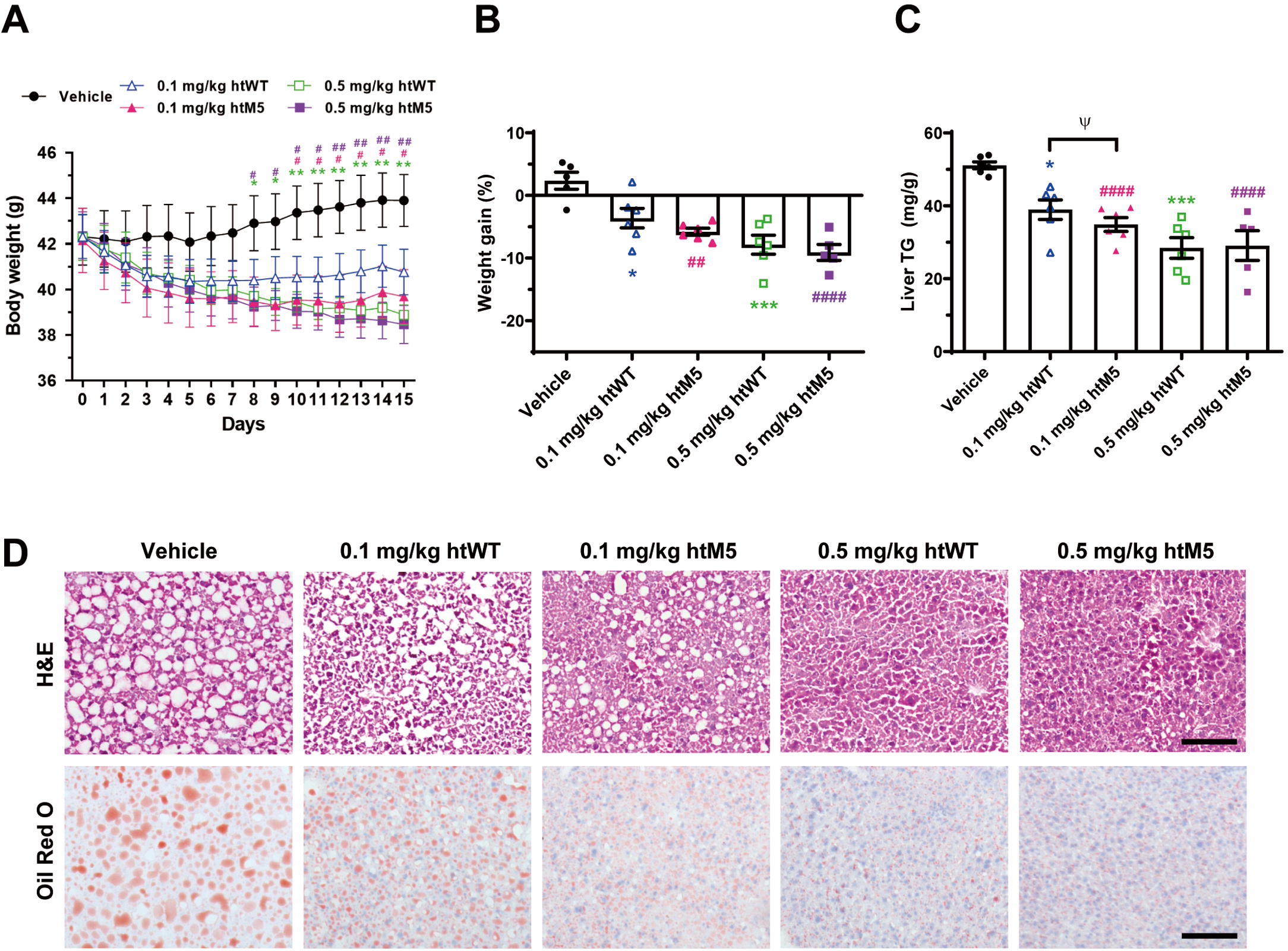
*In vivo* activity tests of htM5. HFD-fed mice were injected with the wild type htFGF21 (htWT) or htM5 intraperitoneally at doses of 0.1 or 0.5 mg/kg once daily for 14 days. *(A)* Changes in body weight throughout the injection. *(B)* Percent weight gains on the following day after the last injection. *(C)* Effects of the injection on liver TG contents. *(D)* H&E and Oil Red O staining of liver sections showing better improvement in htM5 than htWT for the vacuole and lipid accumulation. Data are represented as mean ± SEM, calculated using two-way ANOVA followed by Tukey’s multiple comparisons test, *, #, and Ψ *p* < 0.05, ** and ## *p* < 0.01, *** *p* < 0.001, #### *p* < 0.0001. n = 5–6 mice/group. *, **, and *** indicate vehicle-injected mice versus the htWT-injected mice. #, ##, and #### indicate vehicle-injected mice versus the htM5-injected mice. Ψ indicates the htWT -injected mice versus the htM5-injected mice. Scale bars: 100 μ (H&E), 50 μ (Oil Red O).

Although the vacuolization was still observed in the both livers of mice injected with 0.1 mg/kg of the htFGF21 or htM5, the degree of vacuolization was reduced compared with vehicle-injected mice and the effect of htM5 was greater than that of htFGF21 (Fig. 7*D*). Similarly, Oil red O staining revealed that either the htFGF21 or htM5 administration suppressed HFD-induced hepatic lipid accumulation. htM5 inhibited lipid accumulation at a higher degree than htFGF21 at the dose of 0.1 mg/kg (Fig. 7*D*). Taken together, these results suggest that the introduction of five point-mutations gave rise to the htM5 mutant that is likely to exert better effects for ameliorating NAFLD than the wild-type.

## Conclusion

Characterization of protein stability is an integral step in biopharmaceutical development (39) and the monitoring of thermal behavior is a classical probe to investigate protein stability (40). During examining temperature-dependent structural changes of FGF21, we discovered its temperature-responsive structural reversibility. In addition, we identified the β10-β12 loop to be associated with this remarkable thermal behavior based on the fact that the loop replacement only perturbed thermal response of FGF21 without affecting structural and functional properties. These ever-first findings led us to employ heating as an efficient purification step to separate recombinant FGF21 from heat-labile proteins. Heating has another advantage to prevent the proteolytic cleavage of the essential but fragile C-terminal loop of FGF21 since heat-labile host proteases lose their activities due to thermal denaturation. Consequently, heating-based strategy can be widely applied for the purification of FGF21 analogs that are under therapeutic development.

Five point-mutations (S195G, P199G, S200G, H201Q, and A208T) were given to the C-terminal loop to design FGF21 variants with improved therapeutic potential. S195G, S200G, H201Q, and A208T are firstly reported in this study whereas P199G is a well-known mutation to escape the proteolytic attack by a serum protease (24). H201Q and A208T were designed based on sequence comparison (Fig. S7), indicating that amino acid substitutions at non-conserved sequence positions among FGF21 orthologs can be used to design FGF21 variants. The side chains of Ser195 and Ser200 have no interactions with β-klotho, an indication of little contribution of their side chains to β-klotho binding. Therefore, the glycine introduction instead of those residues seems to be beneficial for co-receptor binding due to the increased flexibility of the C-terminal loop.

## Experimental procedures

### Cloning, expression, and heat-treatment-mediated purification of human FGF21

The coding genes of the wild type FGF21 (residues 33–209), the β-trefoil region of FGF21 (residues 41–173; core-FGF21), and all FGF21 variants (FGF21-LW, core-FGF21- LW, M2, M3, and M5; see the “Results and discussion” section) were synthesized using a TEV protease cleavage site (ENLYFQ/G) at the N-terminus (COSMOGENETECH, Republic of Korea). The synthesized genes were inserted at the *BamH*I and *Sal*I sites of the expression vector pET32a (+) (Novagen, Darmstadt, Germany) to fuse Trx to the N-terminus of the target proteins. The resulting constructs were transformed into *Escherichia coli* strain Rosetta (DE3). The transformants were grown to an OD_600_ of 0.6–0.7 in Luria-Bertani (LB) media containing 50 μg/mL ampicillin and chloramphenicol at 37 °C, and 0.5 mM isopropyl-D-thiogalactoside (IPTG) was added for the expression of the Trx-fused target proteins.

After 42 h of incubation at 15 °C, cells were harvested using centrifugation, resuspended in a buffer (10 mM Tris pH 8.0 and 200 mM NaCl), and disrupted using sonication. To remove cellular debris, centrifugation was performed at 10,000 x g for 40 min at 4 °C. The supernatant was heated at 100 °C for 10 min in a water bath and subsequently centrifuged to secure only soluble fractions. The heat-treated soluble fractions were loaded into a nickel-nitrilotriacetic acid column (Cytiva, Marlborough, MA, USA) and incubated with 0.3 mg/mL

TEV protease overnight at 22 °C to detach target proteins from the Trx tag bound to the resin. The unbound fractions containing target proteins were applied to the Q Sepharose^®^ Fast Flow column (Cytiva) equilibrated with A buffer (10 mM Tris pH 8.0). The elution was performed using a linear gradient of 0–1.0 M NaCl in A buffer. The fractions containing target proteins were collected and finally loaded onto a HiLoad^®^ 16/600 Superdex^®^ 75 pg column (Cytiva) equilibrated with a buffer consisting of 10 mM NaH_2_PO_4_ (pH 8.0) and 200 mM NaCl. Purification was performed by using the AKTA-FPLC system (Cytiva). Proteins used for CD, DLS, and spectrophotometry were purified without heat treatment. Commercial FGF21 (cat# CYT-474) was purchased from ProSpec, Israel.

### Cloning, expression, and purification of human FGF2

The coding gene of FGF2 (residues 143–288) was synthesized at the *Nde*I and *Xho*I sites of the expression vector pET17b (+) (Novagen). The resulting construct was transformed into *Escherichia coli* strain Rosetta (DE3) pLysS. The transformant was grown to an OD_600_ of –0.7 in LB media containing 50 μ /mL ampicillin and chloramphenicol at 37 °C, and 0.5g mM IPTG was added.

After 20 h of incubation at 20 °C, cells were harvested using centrifugation, resuspended in a buffer (10 mM Tris pH 8.0 and 200 mM NaCl), and disrupted using sonication. To remove cellular debris, centrifugation was performed at 10,000 x g for 40 min at 4 °C. The soluble fractions were loaded into a heparin column (Cytiva) and eluted using a linear gradient of 0.5–2.0 M NaCl in 10 mM Tris pH 8.0 buffer. The fractions containing target proteins were collected and finally loaded onto a HiLoad^®^ 16/600 Superdex^®^ 75 pg column (Cytiva) equilibrated with a buffer consisting of 10 mM NaH_2_PO_4_ (pH 8.0) and 200 mM NaCl. Purification was performed using the AKTA-FPLC system (Cytiva).

### Circular dichroism spectroscopy

CD experiments were performed using a Jasco J-1500 equipped with Jasco PTC-517 Peltier cell holder, a thermostat (Jasco Corporation, Tokyo, Japan) and bath circulator (RW3-2025; JEIO TECH Co., Daejeon, Korea). CD spectra were collected with a scanning speed of 50 nm/min, a digital integration time of 4 s, and a bandwidth of 1 nm. Quartz cuvettes with a path length of 1 mm were employed. The signal of a protein-free buffer containing 10 mM NaH_2_PO_4_ (pH 8.0) and 20 mM NaCl was subtracted from all CD spectra. Each spectrum, an average of 5 scans, was normalized to molar ellipticity (θ) using the mean weight residue and concentration by using a program provided by the manufacturer.

To monitor the thermal denaturation, CD spectra of 0.2 mg/mL proteins were collected every 1 °C at 215 nm as the temperature was changed from 20 °C to 100 °C with a heating rate of 1 °C/min. Spectral curves were fitted using the OriginPro 2021 program (Origin Lab Corporation, Northampton, MA, USA) through a non-linear adjustment by the Boltzmann method. When temperature reached 20, 40, 60, 80, and 100 °C during the thermal denaturation experiments, the far-UV CD spectra were obtained in the range of 190–260 nm. The near-UV CD spectra were obtained in the range of 240–340 nm using 5 mg/mL of proteins. Heating rate was set at 5 °C/min and the temperature was maintained for 10 min at 20, 40, 60, 80, and 100 °C. The far- and near-UV CD spectra were collected by continuous scanning at 0.5 nm intervals. The structural reversibility was checked by stepwise cooling of the protein solution from 100 °C to 20 °C.

### Turbidity measurement

The optical density at a wavelength of 600 nm was measured at 75 °C using the V-730 UV-Vis Spectrophotometer (Jasco Corporation) equipped with a bath circulator (DH.WCH00408; DAIHAN SCIENTIFIC Co., Gangwon-do, Korea). The concentration of proteins in a buffer consisting of 10 mM NaH_2_PO_4_ (pH 8.0) and 200 mM NaCl was 25 mg/mL. The value of optical density was determined by subtracting the signal of the protein-free buffer. Measurements began 15 s after putting samples into the spectrophotometer.

### Dynamic light scattering

DLS experiments were conducted at 20 °C using a Zetasizer Nano ZS apparatus (Malvern Instruments Ltd., Malvern, UK) by setting an automatic attenuator. Proteins having a concentration of 25 mg/mL in a buffer consisting of 10 mM NaH_2_PO_4_ (pH 8.0) and 200 mM NaCl were centrifuged at 13,000 rpm for 15 min at 4 °C. The supernatant (1 mL) was transferred to a glass cuvette having a path length of 1 cm that was located at a position of 4.65 mm. The light scattering intensities were collected 14 times at an angle of 173° using a 10 s acquisition time. The size distributions of the target proteins were calculated using the correlation functions by employing the “general purpose mode” in a Zetasizer software v7.13 (Malvern Instruments Ltd.).

### FGFR1c activity assay

To determine the activation level of FGFR1c by FGF21, we used the iLite^®^ FGF-21 assay ready cells (cat# BM3071; SVAR, Malmo, Sweden). These are HEK293 cells modified to express FGFR1c, β-Klotho, GAL4 DNA binding domain fused trans-activation domain of Elk1, GAL4 DNA binding sequence contained reporter gene (firefly luciferase), and renilla luciferase. Cells were seeded with a different concentration of FGF21 proteins in 96-well plates and incubated at 37 °C in a humidified 5% CO_2_ incubator. After 6 h, luciferase activity was determined using the Dual-Glo luciferase assay system (E2920; Promega, Madison, WI, USA) and GloMax^®^96 microplate luminometer (Promega).

### Cell culture and glucose uptake assay

3T3L1 preadipocytes were incubated in Dulbecco’s modified Eagle’s medium (DMEM; GIBCO, Life Technologies Ltd., Paisley, UK) supplemented with 10% bovine serum (GIBCO) and 1% penicillin/streptomycin (GIBCO) at 37 °C in a humidified 5% CO2 incubator. Two days after cells had reached confluence, adipogenesis was stimulated. Cells were maintained in DMEM containing 10% fetal bovine serum (FBS; GIBCO), 0.5 mM IBMX, 0.25 μM DEX, and 5 μg/mL insulin for 2 days. Next, cells were incubated in DMEM containing 10% FBS and 1 μg/mL insulin. After 2 days, cells were maintained in DMEM containing 10% FBS for 4 days. Cellular glucose uptake was determined using the Glucose Uptake-Glo^TM^ Assay (J1342; Promega). Briefly, cells were incubated with FGF21 proteins for 6 h in growth media, followed by incubation in glucose- and serum-free DMEM for 30 min. Cells were treated with 2-deoxyglucose for 10 min. Next, we measured the level of accumulated 2-deoxyglucose-6-phosphate (2DG6P) using the assay kit according to the manufacturer’s protocol using a GloMax^®^96 microplate luminometer (Promega).

### Western blot analysis

To determine FGFR signaling, 3T3-L1 cells were incubated in serum-free DMEM. After 1 h, cells were treated with FGF21 proteins for 15 min. Protein samples were prepared using lysis buffer containing 20 mM Tris-HCl (pH 7.4), 1% NP-40, 10 mM Na_4_P_2_O_7_, 100 mM NaF, 2 mM Na_3_VO_4_, 5 mM EDTA, and a protease inhibitor cocktail (#87786; ThermoFisher Scientific, MA, USA). Whole cell lysates (20 µg) were subjected to SDS-PAGE and immunoblotted with specific antibodies against pFRS2a (#3864; Cell signaling Technology, Beverly, MA, USA), FRS2 (sc-17841; Santa Cruz Biotech, Santa Cruz, CA, USA), pERK (#4370; Cell signaling Technology), and ERK (#4696; Cell signaling Technology). Blots were developed using Clarity™ Western ECL Blotting Substrates (Bio-Rad, Hercules, CA, USA), ECL-Prime (ThermoFisher Scientific), or EZ-Western Lumi Femto (DoGen, Seoul, Korea) with a ChemiDoc Imaging System (Bio-Rad)

### Animals and treatments

All animal experiments were conducted in accordance with the guidelines on animal care and use as approved by the Institutional Animal Care and Use Committee of Daegu Gyeongbuk Institute of Science and Technology (DGIST-IACUC-19121001-0001). Male C57BL/6 mice at 7 weeks of age were purchased from Koatech. Mice were housed under a 12 h light/12 h dark cycle (lights on from 7:00 am to 7:00 pm) in individually ventilated cages (1 mouse per cage) with chip bedding at 23 °C ± 3 °C and a relative humidity of 50% ± 10%. Before the HFD experiments, mice were acclimated to the housing facility for 1 week and given ad libitum access to water and NCD (12% kcal from fat; Lab Supply, 5053). After 10 weeks of a NCD or HFD (60% kcal from fat; Envigo, 06414) treatment, HFD-fed mice were randomly assigned to vehicle or FGF21 treatment groups. The randomization was stratified by body weight. The NCD-fed mice were intraperitoneally injected with vehicle (DPBS, 21-031-CV; Corning, NY, USA) and the HFD-fed mice were intraperitoneally injected with vehicle, wild type FGF21, or mutant FGF21 once daily for 2 weeks. All mice had free access to water and food while remaining on their respective diets during the drug treatment periods. The single injection was performed between 4:00 pm and 6:00 pm, and body weight and food consumption were measured immediately before each injection every day. The following day after the last injection, mice were sacrificed to collect livers after noting final measurements.

### Liver TG quantification

Hepatic TG content was measured using Triglyceride Quantification Colorimetric/Fluorometric Kit (K622-100; BioVision, Milpitas, CA, USA). Livers were homogenized in 1 mL of a solution containing 5% NP-40 in water, heated to 80–100 °C, and allowed to cool to room temperature; this step was repeated twice. Insoluble material was removed using centrifugation at 16000 x g for 2 min. Extracted TG was diluted six-fold with distilled water. TG level was measured according to the manufacturer’s instructions. Individual TG levels were normalized to liver weight.

### Liver histology

Histological analyses of the NAFLD tissues were performed using the H&E Stain Kit (Hematoxylin and Eosin) (ab245880; Abcam, Cambridge, UK) and the Oil Red O Stain Kit (Lipid Stain) (ab150678; Abcam). Each staining was performed according to the manufacturers’ protocols with minor modifications. Livers were rapidly excised from mice, embedded in FSC 22 Frozen Section Media (3801480; Lecia Biosystems, Nussloch, Germany), and frozen on dry ice. Next, fresh frozen sections were obtained on a cryostat (CM3050S; Lecia Biosystems) and fixed using ice cold 4% paraformaldehyde solution (p2031; Biosesang, Seongnam, Korea) for 15 min. The sections were cut into 5 and 7 μm sections for H & E and Oil Red O staining, respectively.

### Data availability

Data are available in the article or Supplementary Information. Source data are provided with this paper.

## Supporting information

Supprting Information

## Supporting information

This article contains supporting information.

## Author contributions

Y.-E.J., K.W.L., and J.H.C. Methodology, Formal analysis, Writing - original draft, review & editing; D.-W.B., B.-G.J., Y.-J.J., Y.J.A., and K.K. Methodology, Writing - original draft; G.S.L., L.-W.K., and J.H.M. Methodology; J.-H.L., E.-K.K., H.-S.Y., and S.-S.C. Conceptualization, Supervision, Investigation, Funding acquisition, Resources, Methodology, Writing - original draft, review & editing.

## Funding and additional information

This work was supported by the project entitled “Development of Biomedical materials based on marine proteins” (the Ministry of Oceans and Fisheries, Republic of Korea).

## Abbreviations

The abbreviations used are: FGF, Fibroblast growth factor; FGFR, FGF receptor; CD, circular dichroism; HFD, high-fat diet; NCD, normal chow die; HSPG, heparin/heparan sulfate proteoglycan; TG, triglycerides; NAFLD, non-alcoholic fatty liver disease; DLS, dynamic light scattering; FRS2, FGFR substrate 2; ERK, extracellular signal-regulated kinase; FAP, Fibroblast activation protein; LB, Luria-Bertani; IPTG, isopropyl-D-thiogalactoside; H&E, hematoxylin and eosin.

