## Supplementary material for "Temperature-responsive structural reversibility of FGF21 and structure-based design of its variant with enhanced potency": Supprting Information

Table of contents

**Supporting Information**	3

Figure S1. Observation of turbidity	3

Figure S2. Structure-based sequence alignment among human FGFs	4

Figure S3. Time-dependent plotting of OD600 at 75 ºC	5

Figure S4. SDS-PAGE analysis of the Trx-FGF21 before (-) and after (+) heating	6

Figure S5. DLS analysis of nFGF21 and htFGF21.	7

Figure S6. The structure of the C-terminal loop of human FGF21 in complex with β-klotho	8

Figure S7. Sequence alignment of the C-terminal loops (residues 193-209 in FGF21) of mammalian FGF21s	9

**Supporting Information**

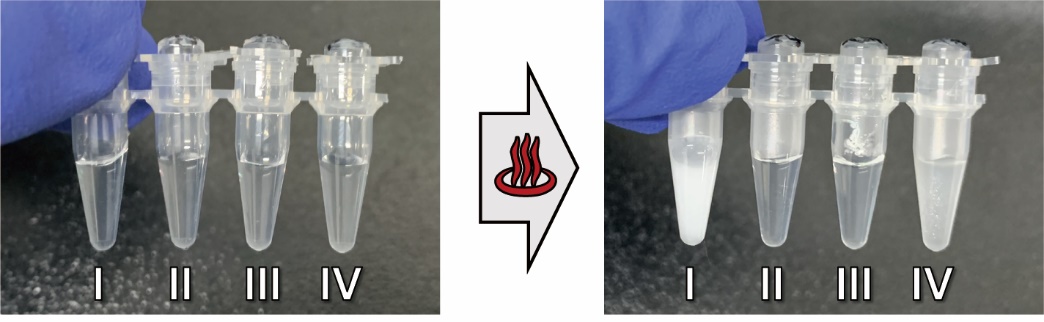

**Figure S1. Observation of turbidity.** The photos show the turbidity of protein solutions in 0.2 ml tubes before (left) and after (right) heating at 100℃. I, II, III, and IV represent FGF2, the wild type FGF21, core-FGF21, and core-FGF21-LW, respectively.

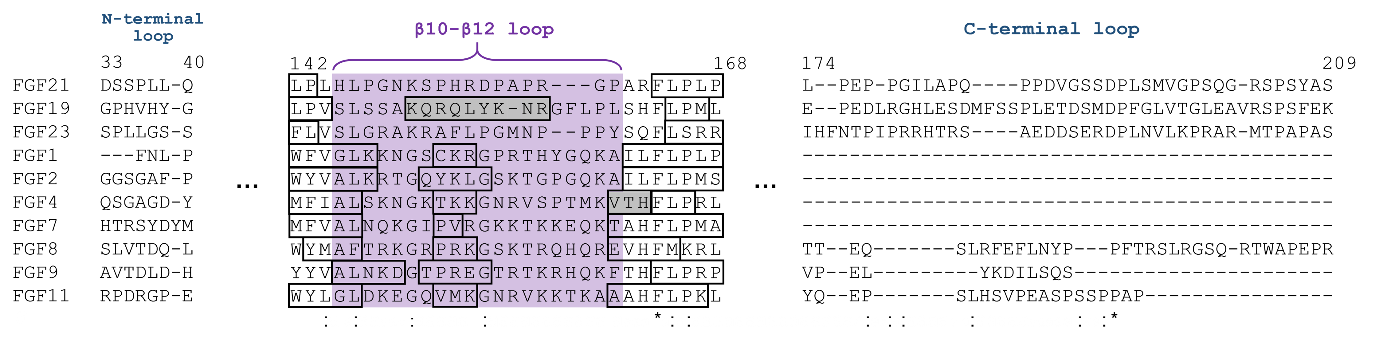

**Figure S2. Structure-based sequence alignment among human FGFs.** Gray boxes indicate α-helices and empty boxes do β-strands. The region corresponding to the β10-β12 loop of FGF21 is shaded in violet. Dashes represent gaps introduced to optimize the alignment. The asterisks and colons indicate identical amino acid residues and conserved substitution, respectively.

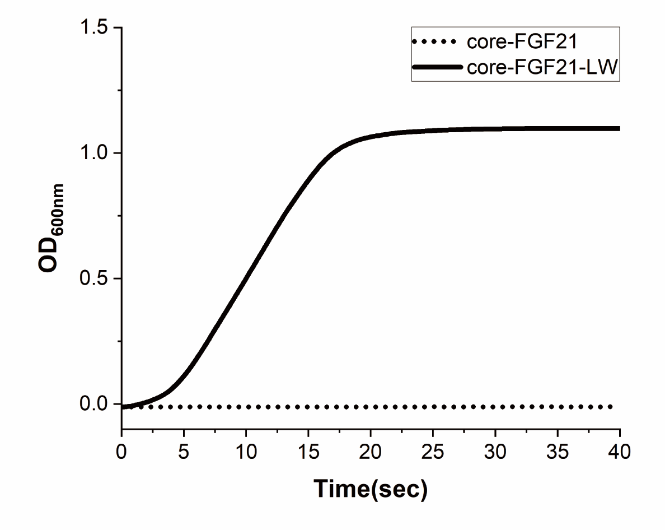

**Figure S3.** **Time-dependent plotting of OD_600_ at 75 ºC.**

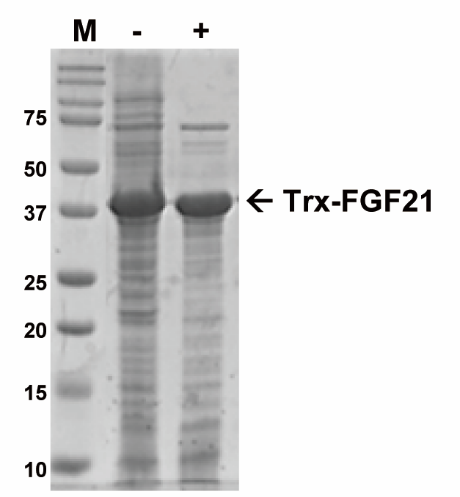

**Figure S4. SDS-PAGE analysis of the Trx-FGF21 before (-) and after (+) heating.** M represents protein markers whose sizes are given in kDa on the left. Arrow points out the position of target protein.

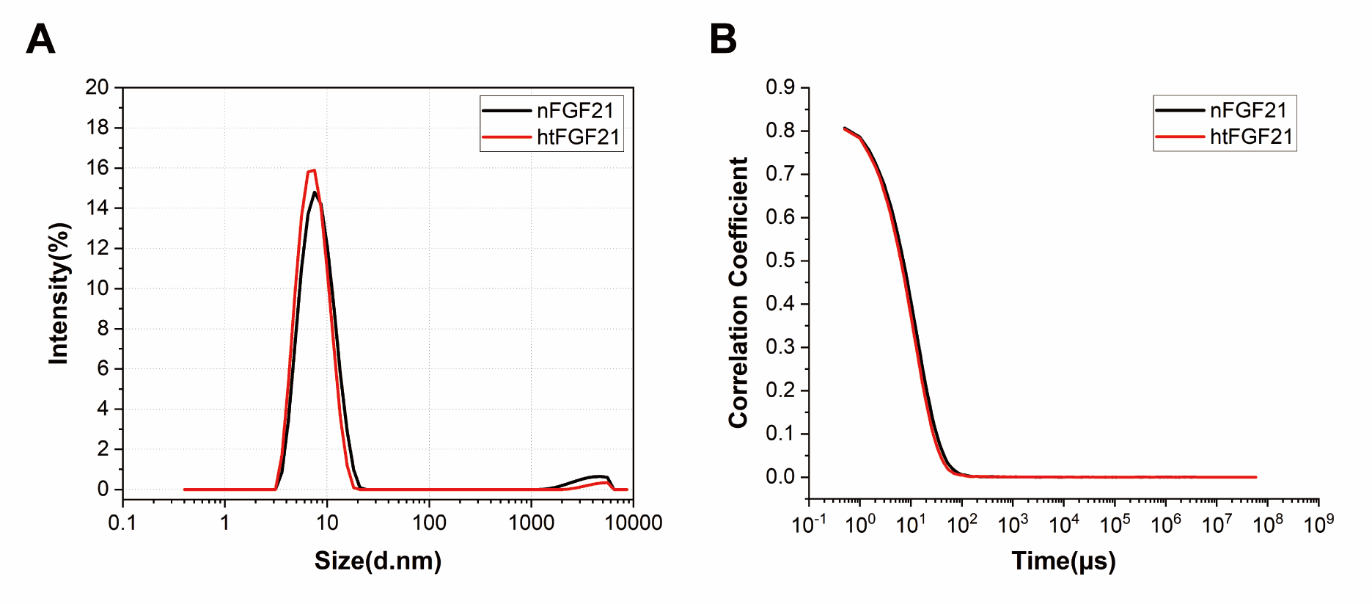

**Figure S5. DLS analysis of nFGF21 and htFGF21.** (A) Comparison of the size distribution histograms between nFGF21 and htFGF21. (B) Raw data of autocorrelation functions detected by DLS. FGF21 purified through heat treatment is represented by htFGF21 while nFGF21 refers to FGF21 purified without heating.

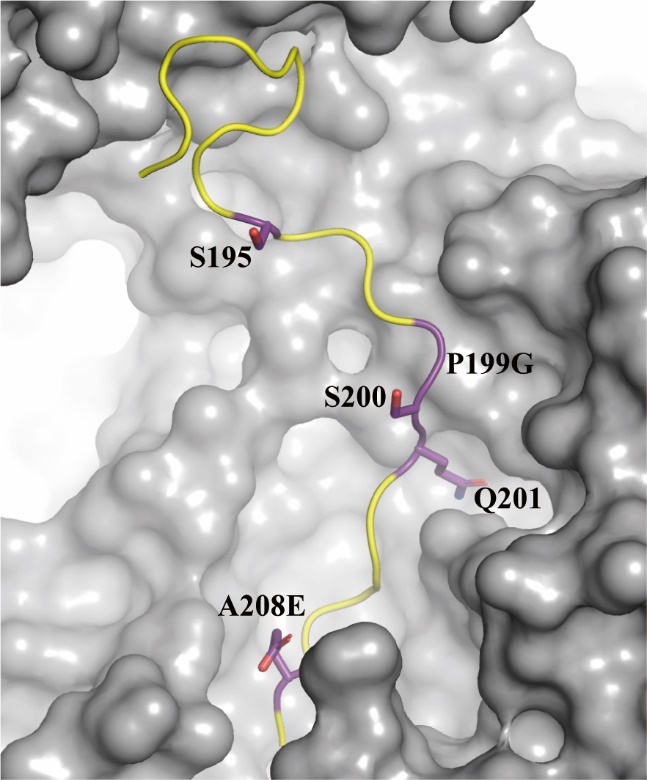

**Figure S6. The structure of the C-terminal loop of human FGF21 in complex with β-klotho (PDB code: 5VAQ) in which the C-terminal loop contains P199G and A208E mutations.** The C-terminal loop is shown in yellow cartoon with five mutation sites in purple sticks. β-Klotho is shown as surface representation. Nitrogen and oxygen atoms in sticks are colored in blue and red, respectively.

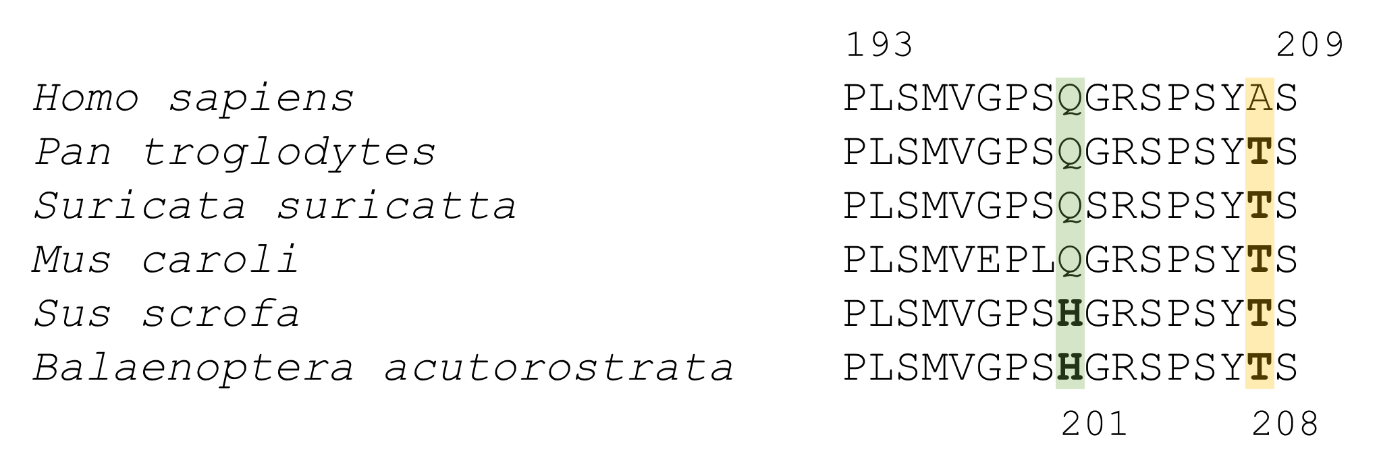

**Figure S7.** Sequence alignment of the C-terminal loops (residues 193-209 in FGF21) of mammalian FGF21s. Amino acids at positions 201 and 208 are colored in green and yellow, respectively. Histidine and threonine at positions 201 and 208 are shown in bold. The GenBank accession numbers are as follows: NP_061986.1 (*Homo sapiens*), XP_016791946.1 (*Pan troglodytes*), XP_029780315.1 (*Suricata suricatta*), XP_021024694.1 (*Mus caroli*), NP_001156882.1 (*Sus scrofa*), XP_007168312.1 (*Balaenoptera acutorostrata scammoni*).
